# Deep-layer cortical tracking abruptly collapses in the absence of language comprehension

**DOI:** 10.64898/2026.09.04.747128

**Authors:** ChenTianyi Yang, Andrew Thwaites, Cai Wingfield, Chao Zhang, Alexandra Woolgar

**Author notes:** Contributing authors. These authors contributed equally to this work (ChenTianyi Yang, Andrew Thwaites).

## Abstract

The brain builds meaning from speech in stages, transforming acoustic input into linguistic comprehension. Yet where comprehension separates from general acoustic processing has been difficult to localize, because the two are tightly entangled in continuous speech. Here we align the activity of 145,000 individual artificial neurons of an audio large language model with high-resolution magnetoencephalography, using single-neuron interpretability to track, layer by layer, which computations the cortex follows during natural listening. Contrasting native listeners with listeners hearing an unfamiliar language, while holding the acoustics physically identical, we find that comprehension selectively sustains brain–model alignment through the model’s deep layers. Without comprehension, alignment in the acoustic encoder is preserved but deep-layer tracking abruptly collapses, and the sparse alignments that survive correspond only to physical acoustic features. This pinpoints the stage at which comprehension departs from perception and yields an interpretable, non-invasive index of whether speech is understood.

## 1 Introduction

Speech comprehension unfolds through a hierarchy of processing stages [1, 2]. A central question in the neuroscience of language is where, along this hierarchy, language-specific comprehension diverges from general acoustic processing. In natural, continuous speech this boundary is difficult to localize, because acoustic and linguistic information are tightly entangled: cortical activity that follows the speech signal can be driven strongly by its acoustics alone, and therefore cannot by itself establish that the signal has been understood [3, 4].

Established neural markers of language processing were not designed to resolve this boundary accurately. Markers such as the N400 [5, 6] and multivariate decoding of linguistic states [7, 8] were designed for controlled contrasts or deviance paradigms rather than continuous, task-free listening, and indicate whether comprehension occurred rather than where it departs from acoustic processing. Deep-network models of speech and language now predict cortical responses directly from natural speech [9–12], offering a computational lens on the hierarchy, but the same entanglement persists: model representations mix acoustic and linguistic information, and alignments to the brain are typically obtained by fitting a mapping that pools across many model units or sensors [13, 14], obscuring the contribution of individual computations. Pooling obscures the contribution of individual computations, and multivariate mappings can further capitalize on low-level confounds, potentially inflating apparent alignment [15]. What has been missing is a way to read out, unit by unit, which specific computation the cortex tracks at each moment of natural listening.

Here we align the activity of ∼145,000 individual artificial neurons of a multilingual audio large language model (aLLM) directly with high-resolution magnetoencephalography (MEG) recorded during natural listening. Rather than fitting a high-dimensional mapping, we test each artificial neuron against each MEG sensor with a direct, univariate temporal correlation. This preserves the latency of individual processing streams and permits single-neuron mechanistic interpretability, allowing us to identify which computation each aligned unit performs and at what cortical latency. To isolate the comprehension–perception boundary, we contrast native listeners with listeners hearing an unfamiliar language, holding the acoustic stimulus physically identical while varying only whether it is understood — a well-established paradigm for probing the linguistic hierarchy [16, 17].

We find that in native listeners, cortical tracking follows the model’s representations across its full computational depth, from the acoustic encoder into the deep decoder layers where higher-order linguistic representations emerge. Comprehension selectively sustains this deep-layer alignment: when the same speech is not understood, encoder-stage tracking is preserved but deep-layer tracking abruptly collapses, and the sparse alignments that survive correspond only to physical acoustic features. This localizes the divergence between acoustic perception and language-specific comprehension to a specific computational stage, and establishes a layer-by-layer computational reference for the healthy sound-to-meaning pathway. Because the boundary is defined on natural, continuous speech without an overt behavioral task, it also offers an interpretable, non-invasive index of whether speech is understood, with potential relevance to assessing receptive language where behavioral report is absent or unreliable [18, 19].

## 2 Results

### 2.1 Hierarchical cortical tracking maps the continuous trajectory of natural speech

To establish the baseline trajectory of natural speech processing, we aligned the activity of ∼145,000 individual artificial neurons from the multilingual Qwen2.5-Omni model [20] with MEG recordings of native Russian and English listeners processing continuous, spoken-word narratives in their respective languages (Fig. 1A). Within this model architecture, audio is processed sequentially through three functionally distinct stages: the initial tokenization of frequency-banded audio energy, an ‘encoder’ block that processes the raw acoustic input, and a deeper ‘decoder’ block in which higher-order representations emerge.

**Fig. 1.**
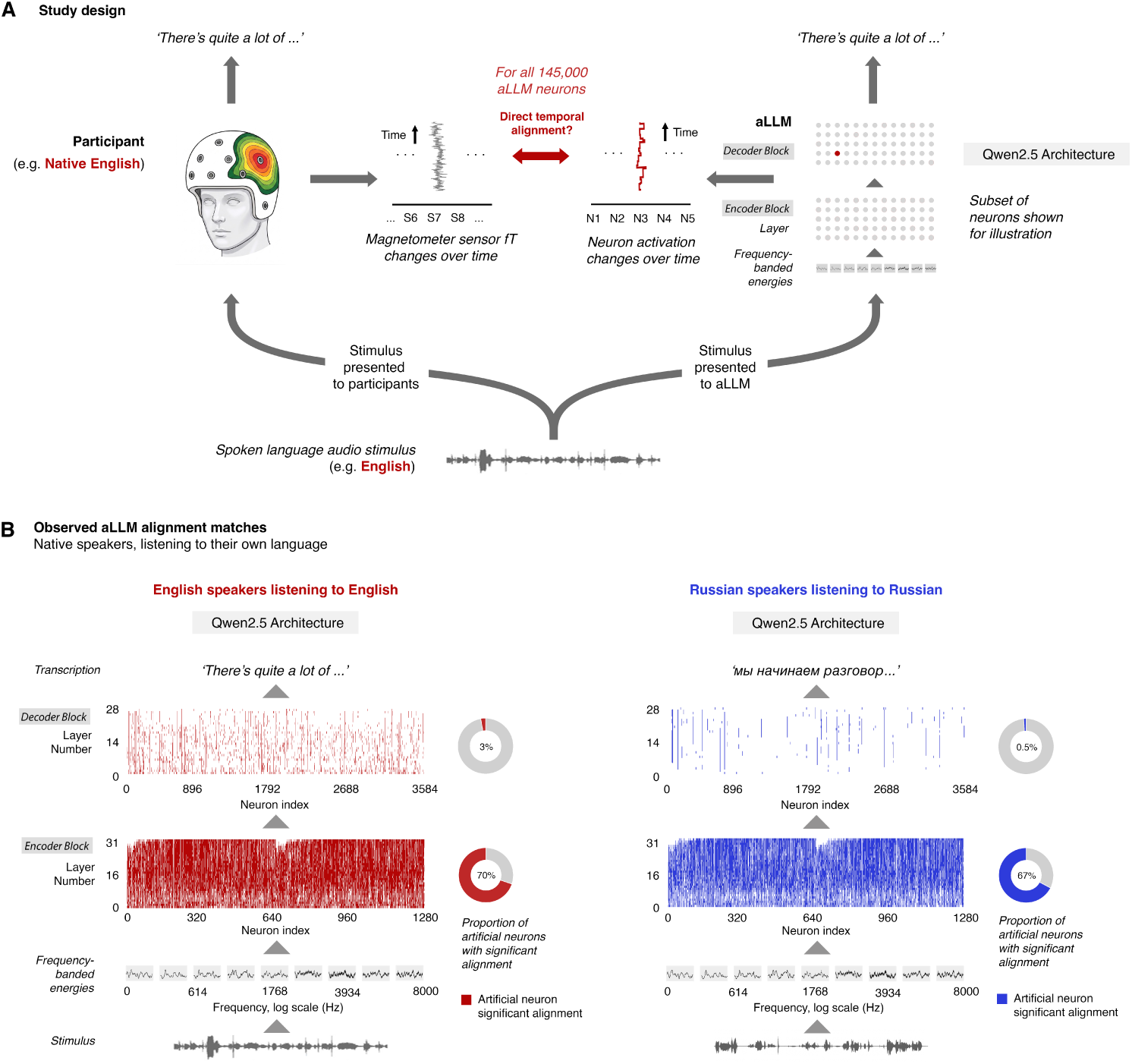
Alignment of individual aLLM neurons to cortical activity during speech comprehension. **(A)** Study design. A continuous natural-speech stimulus is presented both to a native speaker of the language, whose cortical responses are recorded with MEG, and to an audio large language model (aLLM). The activation time course of each individual artificial neuron is then tested for direct temporal alignment against every MEG sensor. **(B)** Significant alignments across the 40,960 encoder units (32 layers × 1,280 dimensions) and 103,936 decoder units (29 layers × 3,584 dimensions). Left (red): native English speakers listening to English. Right (blue): native Russian speakers listening to Russian. Markers indicate units significantly aligned to at least one MEG channel. Structure visible in these index-ordered maps — a bipartite repeat within the encoder and vertical banding across decoder layers — arises from the model architecture rather than the alignment, and is characterized in Supplementary Information (SI1, SI2).

We observed direct temporal alignment between brain and machine in both languages, with a far higher proportion of matches for artificial neurons from the encoder, relative to decoder, layers (native English: encoder neurons 70%, decoder neurons 3%; native Russian: encoder neurons 67%, decoder neurons 0.5%) (Fig. 1B). Representations propagating through successive layers demonstrated an unbroken progression of cortical tracking across the full computational depth of the model, aligning with cortical activity at progressively longer latencies. In the acoustic–phonetic encoder block, this alignment followed a strictly linear trajectory (English: 86.4% variance explained, *p*-value 1.58 × 10*^−^*^14^, ΔBIC(linear *>* quadratic) = 1.38; Russian: 82.7% variance explained, *p*-value 5.82 × 10*^−^*^13^, ΔBIC = 1.73, see Methods, SI4 for details), while alignment in the deeper decoder block progressed with a decelerating, quadratic trajectory (English, 91.3% variance explained, *p*-value 9.17 × 10*^−^*^16^, ΔBIC(quadratic *>* linear) = 14.60; Russian, 32.2% variance explained, *p*-value 1.63×10*^−^*^3^, ΔBIC = 3.85) (Fig. 2A left).

**Fig. 2.**
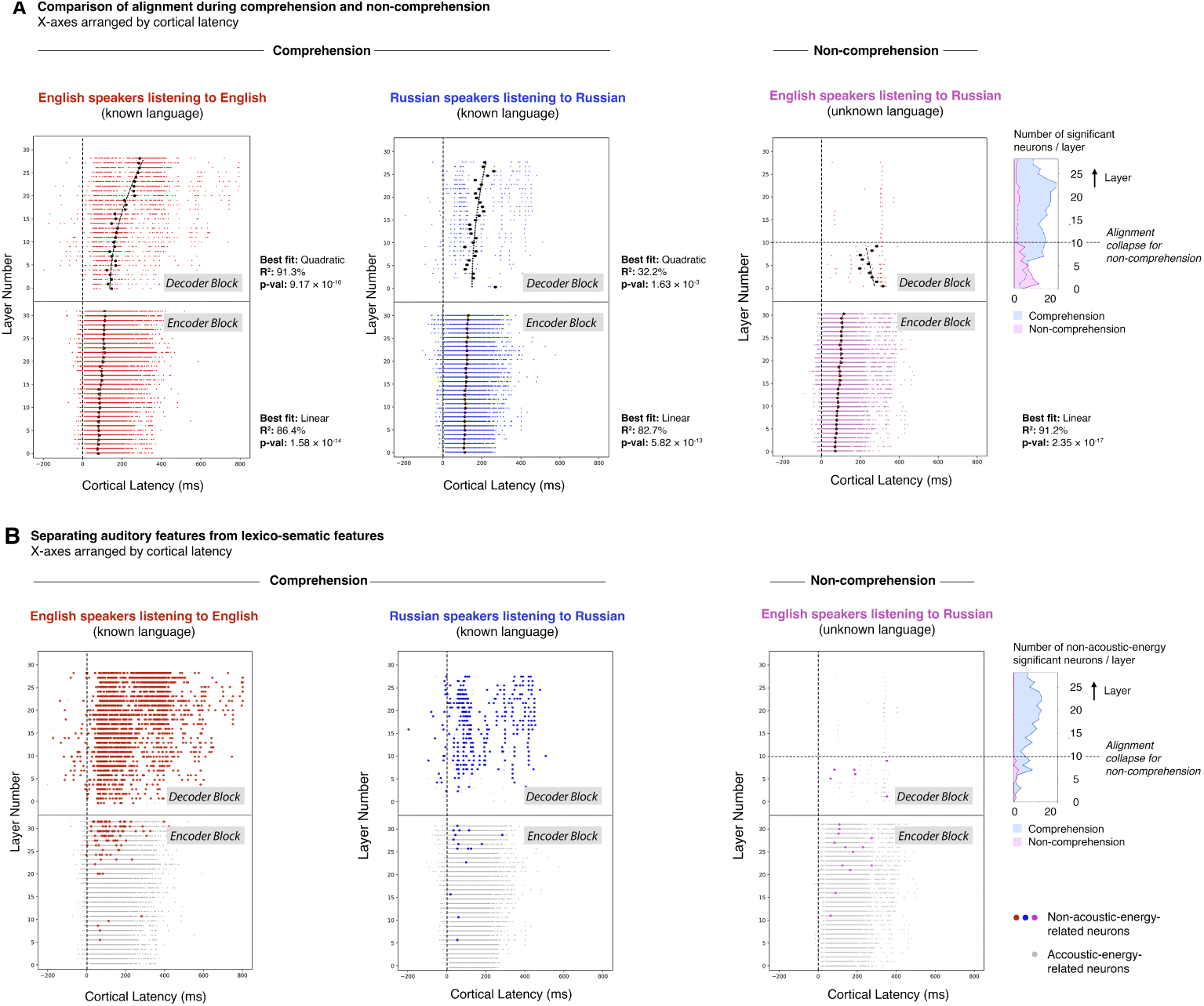
Alignment of aLLM neurons during comprehension and non-comprehension. **(A)** Left two panels (red and blue): comprehension conditions: same results as Fig. 1B, but with the *x*-axis representing the cortical latency (ms) at which the artificial neuron’s response was expressed in the MEG data. Black points are the mean latency of all significant neurons in the layer, with linear and quadratic fits applied to encoder and decoder blocks, respectively (see SI4). Right panel (purple): uncomprehended speech (native English speakers listening to Russian). Collapse boundary marked at layer 10 (4.6). Inset shows the number of significant neurons per layer for both comprehended (blue) and non-comprehended (purple) speech. **(B)** As panel A, with non-acoustic-energy-related neurons highlighted.

Consistent with the known left-lateralisation of higher-order language processing, the peak deep-layer alignments were predominantly left-lateralised, whereas earlier-layer alignments were more bilaterally distributed (Supplementary Information, Fig. SI1).

### 2.2 Deep-layer cortical tracking collapses in the absence of language comprehension

We next sought to functionally isolate the divergence point between language-general and language-specific processing by disrupting the listener’s ability to extract meaning from the acoustic stream. To achieve this, we measured cortical entrainment in a counterfactual cohort of native English, non-Russian-speaking participants who listened to the same Russian audio stimulus, but without comprehending it. In the absence of language comprehension, cortical tracking within the encoder block maintained the same uninterrupted linear trajectory observed in native speakers (91.2% variance explained, *p*-value 2.35 × 10*^−^*^17^). However, the progression of neural tracking collapsed within the deeper decoder block: the number of artificial neurons exhibiting significant tracking with the MEG sensors fell sharply and did not recover across the deeper decoder layers (Fig. 2A right).

To locate this collapse without imposing an arbitrary threshold on the per-layer neuron count, we identified the boundary that best separated the high-tracking early layers from the low-tracking later ones. For each candidate boundary we computed a rank-based separation statistic, *A*(*b*) — the probability that a randomly chosen pre-boundary layer contains more significantly tracking neurons than a randomly chosen post-boundary layer — and selected the boundary that maximised it (See Methods). This placed the boundary at layer 10, with near-total separation between the two regimes (*A* = 0.99, where 1 denotes perfect separation): beyond layer 10, high neuron counts did not recur. A permutation test correcting for the search over candidate boundaries confirmed that separation this clean was unlikely to arise by chance (*p* = 0.003). The boundary’s location was well constrained, though not pinned to a single layer: a conservative 90% confidence set spanned layers 6–11, and it was sharply identified only under the assumption of an abrupt rather than gradual transition (Fig. 3; Methods).

**Fig. 3.**
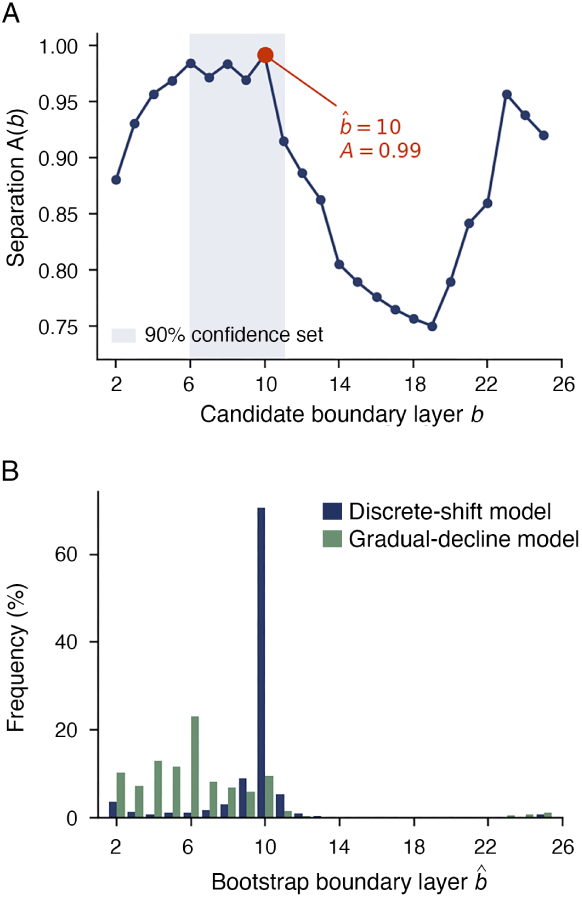
The collapse of deep-layer tracking is localized to a specific layer, computed from the per-layer counts in Fig. 2A. **(A)** Separation statistic *A*(*b*) across candidate boundaries, maximized at layer 10 (highlighted), beyond which high neuron counts do not recur. Shading marks the 90% confidence set for the boundary location (layers 6–11). Statistic not calculated for starting and trailing layers 0-1 and 27-28. **(B)** Bootstrap distributions of the estimated boundary ^^^*b* under a discrete-shift model and a gradual-decline model. The location is sharply identified only under the discrete-shift assumption.

### 2.3 Surviving deep-layer alignment exclusively tracks physical acoustic features

Given the abrupt collapse of deep-layer tracking, we applied mechanistic interpretability techniques to characterize the sparse subset of early decoder neurons that maintained alignment in uncomprehending listeners. We hypothesized that without active semantic integration, this surviving cortical tracking reflected purely physical, language-invariant properties. Evaluating these activations against established psychophysical models [21], we found that all surviving neurons in deep layers of the uncomprehending cohort were highly correlated with acoustic-specific frequency-band loudnesses (Fig. 2B). Furthermore, of the remaining six *non*-loudness-related decoder neurons aligned during non-comprehended speech, all were entirely explained by a simple word-duration feature (Kruskal–Wallis test, *H >* 36.5, *p <* 0.001, see Methods).

## 3 Discussion

Aligning individual artificial neurons of an audio large language model directly with high-resolution MEG, we localized the computational stage at which language-specific comprehension diverges from language-general perception during natural speech. Comprehension proved necessary to sustain this deep-layer alignment: in its absence, encoder-stage tracking persisted while deep-layer tracking collapsed, and the units that remained aligned tracked only physical acoustic features. Together, this provides a layer-by-layer computational reference for the healthy sound-to-meaning pathway against which departures from comprehension can be measured.

Prior work has established that speech comprehension relies on a hierarchical progression, successfully mapping how neural representations transition from acoustic perception to higher-order linguistic integration [2, 16, 22]. While the stages of this hierarchy have traditionally been delineated by observing how higher-order neural tracking degrades—either as the acoustic signal itself is degraded [23] or when the language is unfamiliar [17, 24]—such approaches reveal that this tracking is reduced, but not the precise computational stage at which language-specific comprehension diverges from language-general perception. Because our design holds the acoustics physically identical while varying only comprehension, the collapse we observe reflects the loss of comprehension rather than any reduction in the sensory signal, and aligning individual artificial neurons to cortical activity localizes this divergence to an exact computational stage. Likewise, recent deep-learning alignments have offered a flexible computational lens on the broader cortical hierarchy [10, 12, 25], but because they pool across many units they cannot attribute alignment to individual computations. Reading out single neurons instead lets us specify which representations survive when comprehension fails, and we find that these correspond exclusively to physical acoustic properties. This suggests that when semantic integration fails, the cortex does not simply disengage but falls back on tracking low-level acoustic structure.

Two boundary conditions define the scope of these claims. The first concerns generality: while the sound-to-meaning trajectory itself replicates across two languages and cohorts, the collapse of deep-layer tracking was established with a single non-comprehension dataset and one continuous stimulus. Whether the same boundary emerges across other languages, stimuli, and models is a directly testable prediction of this account. The second concerns resolution: these analyses rest on group-averaged responses, which establish the phenomenon at the population level but do not yet resolve individual listeners. Locating the boundary reliably in individuals would help distinguish perceptual from higher-order comprehension failures — including in individuals for whom behavioral report is absent or unreliable [18, 19]. Achieving this requires single-subject reliability, attainable in principle through higher-resolution recordings or individualized model tuning, and is a prerequisite for translational application.

A complementary direction concerns the representations unique to comprehension itself. Having characterized the perceptual features that survive in its absence, a natural next step is to use single-neuron interpretability [26, 27] to map the fuller suite of semantic features that distinguish successful comprehension [28]. More broadly, by reading out individual computations directly from natural speech, this approach establishes single-neuron brain–model alignment as a specific and interpretable assay of the human language system.

## 4 Methods

### 4.1 Open datasets

Three datasets were used in the study. Both native speaker datasets are open access, while the non-comprehended ‘Native English speakers listening to Russian’ dataset was recorded specifically for this study, using the same equipment. All three datasets are summarised in Table 1.

**Table 1.** Summary of datasets used in this study.

| Dataset | Stimulus length | Participants | Repetitions |
| --- | --- | --- | --- |
| Kymata Soto English [3] | 6 min 40 sec | 20 | 8 |
| Kymata Soto Russian [3] | 6 min 40 sec | 15 | 4 |
| Uncomprehended speech | 6 min 40 sec | 15 | 4 |

#### Kymata Soto English

The Kymata Soto English dataset [29] contains MEG recordings of 20 adults (11 male, 9 female, mean age = 22 years, range = 18–35), all of whom were native English speakers. All the participants had normal hearing and no hearing-related disorders. During the experiment, participants listened to an English-language audio stimulus: a three-way discussion from a British Broadcasting Corporation (BBC) Radio discussions (2 male speakers, 1 female) on the history of ice cream, edited to a length of approximately 400 seconds (6 minutes and 40 seconds). Nineteen participants listened to 8 repetitions of the stimulus; one participant listened to 4 repetitions. All repetitions were delivered in a single recording session. Participants were asked two novel multiple-choice questions on the content of the stimulus after each repetition, and they responded using a button box.

#### Kymata Soto Russian

The Kymata Soto Russian dataset [29] contains the MEG recordings of 15 adults (7 male, 8 female, mean age = 24 years, range = 18–30), all of whom were native Russian speakers. All the participants had normal hearing and no hearing-related disorders. During the experiment, participants listened to a Russian-language audio stimulus: a three-way discussion (2 male speakers, 1 female) on the history of Colombian coffee, edited to approximately 400 seconds (6 minutes and 40 seconds). Participants were asked two novel multiple-choice questions on the content of the stimulus after each repetition, and they responded using a button box. All participants listened to 4 repetitions of the stimulus, delivered in a single recording session for each participant.

Both studies were preprocessed using a common preprocessing pipeline, detailed in section 4.2.4.

### 4.2 Non-comprehended-speech MEG study

We conducted a MEG study using the same stimulus as the Kymata Soto Russian experiment, but using 15 English-speaking participants with no knowledge of Russian.

#### 4.2.1 Stimulus presentation

We used the same audio stimulus as the Kymata Soto Russian experiment [29]. The stimulus was presented at a sampling rate of 44.1 kHz with 16-bit resolution.

#### 4.2.2 Participants

We recruited fifteen participants (7 male, 8 female, mean age = 22 years, range = 19–27), all of whom were native English speakers with no knowledge of Russian. Twelve participants heard 4 repetitions of the stimulus and three participants heard 8 repetitions. All the participants had normal hearing and no hearing-related disorders.

#### 4.2.3 Data acquisition

We used the same data-acquisition protocol to Yang et al. [29]. Within each recording session, MEG data were recorded for participants, and sampled at 1,000 Hz while they listened to the podcast. The MEG data were recorded using a 306 channel VectorView system (Elekta-Neuromag, Helsinki, Finland) containing 102 identical sensor triplets (two orthogonal planar gradiometers and one magnetometer) in a hemispherical array situated in a light magnetically shielded room.

#### 4.2.4 Data preprocessing

The continuous data were notch filtered for the HPI coil and electrical line signals. Independent Component Analysis (ICA) is used to identify the heart beat, vertical eye movement, and horizontal eye movement artifacts in the signals, and those components are manually removed. We then compensated for the head movement and used signal-space separation [30] for the MEG data. The recording was split into 400 epochs of 1,000 ms duration. Each epoch included the 200 ms from before the epoch onset and 800 ms after the epoch finished (taken from the previous and subsequent epochs) to allow for the testing of different latencies. Activity was averaged across participants.

For the source-level analysis in the Supplementary Information, source-current estimation signals were obtained using minimum norm estimations [31] for the 20,484 cortical regions for analysis related to cortical locations.

### 4.3 aLLM Feature Generation

Audio recordings were segmented into 30-second windows and processed using Qwen2.5-Omni [20] with default parameters. We prompted the models to perform speech recognition, using teacher-forcing to constrain the final output to the ground truth transcription generated by reconciling two independently human-corrected automatic speech recognition outputs. Our analysis focused on both the audio encoder (32 layers, 1280 dimensions) and LLM decoder (29 layers, 3584 dimensions). The output of the token embedding layer right before the first decoder transformer block is considered the first of the 29 decoder layers. We extracted activations from the final fully connected layer of each encoder and decoder Transformer block, treating each dimension as an individual neuron. This yielded a total of 40,960 encoder units (32 layers × 1280 dimensions), and 103,936 decoder units (29 layers × 3584 dimensions) for testing. Since the decoder activations align with the word tokens in the output transcription, we employed Montreal forced alignment (MFA [32]) to align these tokens with the audio timestamps, thereby converting the activations into time-series data.

### 4.4 Testing for Cortical Entrainment

While conventional artificial neural network (ANN)-to-brain encoding frameworks frequently utilize multivariate approaches—such as cross-validated ridge regression—to align high-dimensional model layers with neural data, recent work highlights that such approaches can be highly susceptible to low-level, non-linguistic confounds. Specifically, optimized multivariate transformations can inadvertently capitalize on features like word rate or positional signals, potentially inflating predictivity or obscuring the true underlying relationship between the model and the brain [15, 33].

To systematically circumvent these constraints, we eschew intermediate regression-based fitting in favor of a direct, univariate temporal correlation pipeline. This direct approach is explicitly designed to achieve three primary objectives: (1) eliminate the capacity for a high-dimensional mapping model to artificially inflate predictivity, ensuring that any detected alignment reflects an intrinsic, unrotated relationship between individual artificial units and cortical sensors; (2) preserve the fine-grained temporal dynamics and latencies of individual processing streams, which are heavily masked or distorted by multivariate aggregation; and (3) enable strict single-unit mechanistic interpretability, a prerequisite for pinpointing the exact computational layer at which language-specific tracking collapses.

We assessed cortical entrainment by correlating the generated feature functions with the MEG data, following the methods of Thwaites et al. [34–36]. This approach treats the auditory stimulus as the input for both a proposed mathematical transform (in this case, Qwen neuron activations) and the biological cortical processing. A significant match between the model output and the brain signals suggests that the brain performs operations analogous to the proposed transform.

To account for neural processing delays, we optimized for the best latency between the neural data and the function output. Specifically, for each of the activation profiles in the aLLM, we calculated Pearson’s correlation coefficient against every brain-measurement channel. We generated a null distribution by calculating correlations against random trial-wise permutations of the neural data. Welch’s *t*-test was used to determine the significance of the correlations. Because the latency between the audio onset is unknown, we repeated this analysis at 5 ms intervals over a latency window of -200 ms to 800 ms. This procedure was iterated for every channel, every artificial neuron, and every layer of the aLLM.

We reported our results based on statistical significance. To rigorously control for multiple comparisons (201 latency values, 306 MEG channels, and 144,896 individual neurons), we applied a 5-sigma threshold, corresponding to a *p*-value of approximately 2.88 × 10*^−^*^17^. For the dots shown in Figure 2, each represented an individual neuron with the latency value chosen as the one corresponding to the minimum p-value of the alignment for that neuron.

### 4.5 Modelling latency trends across layers

To characterise how alignment latency progressed through the model, we modelled each layer’s mean latency as a function of its depth, separately for the encoder and decoder blocks. For each layer we averaged the latencies of its significantly aligned neurons, requiring at least four such neurons for a layer to be included so that estimates were not skewed by sparse or spurious matches. We fitted two ordinary least squares models to these per-layer latencies: a linear model (latency ∼ intercept + layer) and a quadratic model (latency ∼ intercept + layer^2^), the latter with its stationary point fixed at layer zero and no linear term, so as to encode a strictly accelerating or decelerating trajectory. To compare the two, we computed the Bayesian information criterion (BIC) for each and took their difference, ΔBIC = BIC_quadratic_ − BIC_linear_, such that positive values favour the linear model and negative values the quadratic; magnitudes were interpreted against standard rule-of-thumb thresholds [37]. Full per-dataset fits and BIC values are given in Supplementary Information (Fig. SI2, Table SI1).

### 4.6 Defining the non-recovery boundary

Let *k_ℓ_* denote the number of artificial neurons significantly tracking the MEG sensors at decoder layer *ℓ* (0 ≤ *ℓ* ≤ 28; significance as defined above). To locate the collapse without imposing an *a priori* threshold on these counts, we identified the layer that best separated the high-tracking early layers from the low-tracking later ones.

#### Separation statistic

For each candidate boundary *b* we partitioned the layers into a pre-boundary set {0*, . . . , b* − 1} and a post-boundary set {*b, . . . ,* 28}, and computed a separation statistic

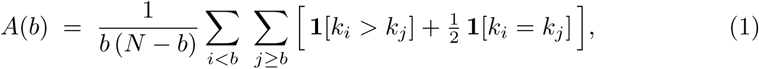

where *N* = 29 is the number of decoder layers. *A*(*b*) is the normalised Mann–Whitney statistic, equivalently a common-language effect size: the probability that a randomly chosen pre-boundary layer has a higher neuron count than a randomly chosen post-boundary layer, with ties counted as one half. Being rank-based, it is insensitive to the magnitude of individual outliers, and is maximised by boundaries after which no large counts recur — the property we take to operationalise collapse without recovery. We estimated the boundary as ^^^*b* = arg max*_b_ A*(*b*) over *b* ∈ {2*, . . . ,* 25}, excluding degenerate partitions.

#### Statistical significance

We assessed whether separation this clean could arise by chance using a permutation test that corrects for the search over boundaries. Under the null hypothesis that the layer counts are exchangeable with respect to layer order, we defined the test statistic *T* = max*_b_ A*(*b*) and recomputed it for 10^5^ random permutations of the counts. Because the maximum is taken over the same set of candidate boundaries in every permutation, the resulting *p*-value accounts for ^^^*b* having been selected from the data and is not inflated by that selection.

#### Boundary uncertainty

Because *A*(*b*) has a broad maximum, we characterised uncertainty in the boundary location with two bootstrap procedures that bracket the assumed form of the transition (10^4^ replicates each, with ^^^*b* re-estimated per replicate as above). The first assumed a discrete regime shift: counts were simulated as Poisson variates with means equal to the empirical pre- and post-boundary counts at ^^^*b* (7.5 and 1.8 neurons, respectively). The second allowed a gradual decline with no step: we fitted a monotone non-increasing profile to the counts by isotonic regression and simulated Poisson counts around the fitted values. We report a conservative 90% confidence set that brackets the distributions from both models, and treat the boundary as sharply identified only under the discrete-shift assumption.

### 4.7 Interpreting aLLM Features

After we obtained the activations from aLLM neurons and identified the ones that are significantly correlated with MEG data, we interpreted those activations as time-series data using time-varying loudness models [38] and the word-duration feature. For the loudness models, we calculated the Pearson’s correlation coefficient between each of the activations of the significant aLLM neurons and the output of the loudness models. We generated a null distribution by calculating correlations against randomly permuted output. Welch’s *t*-test was used to determine the significance of the correlations. We repeated this analysis at 5 ms intervals over a latency window of -500 ms to 500 ms to account for possible delay of encoding loudness information in the aLLM neurons. We applied a 5-sigma threshold of 1.30 × 10*^−^*^10^ for the *p*-value after considering multiple comparisons.

After interpreting the loudness-related aLLM neurons, we obtained the duration of each word piece and tested if it can explain the variation of the corresponding activations. We used the non-parametric Kruskal–Wallis H test [39] to see if the activation distribution is different across the duration groups. The null hypothesis is that the activations for all duration groups come from the same distribution.

## 5 Acknowledgments

This work was supported by UKRI MRC intramural funding to AW (MC_UU_00030/15), a grant from Neuroverse FZCO to AT and CZ, and a National Natural Science Foundation of China (NSFC) grant to CZ (62476151). We would like to thank Ariel Goldstein for feedback on an earlier version of this manuscript.

## 6 Author Contributions

**ChenTianyi Yang**: Formal analysis, Investigation, Software, Visualization, Writing – review & editing. **Andrew Thwaites**: Conceptualisation, Writing – original draft, Software, Visualization. **Cai Wingfield**: Formal analysis, Investigation, Software, Visualization, Writing – review & editing. **Chao Zhang**: Conceptualisation, Writing – review & editing. **Alexandra Woolgar**: Conceptualisation, Writing – review & editing.

## 7 Data Availability

The dataset is available on OSF at https://doi.org/10.17605/OSF.IO/QDZGH.

## 8 Code Availability

The analysis used the Kymata-Core code, available at https://github.com/kymata-atlas/kymata-core.

## Supplementary Information

## Appendix SI1 Analysis of aLLM-brain encoder and decoder block patterns

In the alignment maps plotted by neuron index (Fig. 1B), the decoder block exhibits a spatially uniform distribution of significant alignments with no discernible horizontal pattern in both English and Russian. This absence of structural periodicity reflects the nature of the aLLM architecture: the semantic or functional role of any given artificial neuron within a decoder layer is not computationally constrained to a specific index. Consequently, these higher-order feature representations emerge as arbitrarily indexed networks during model training:

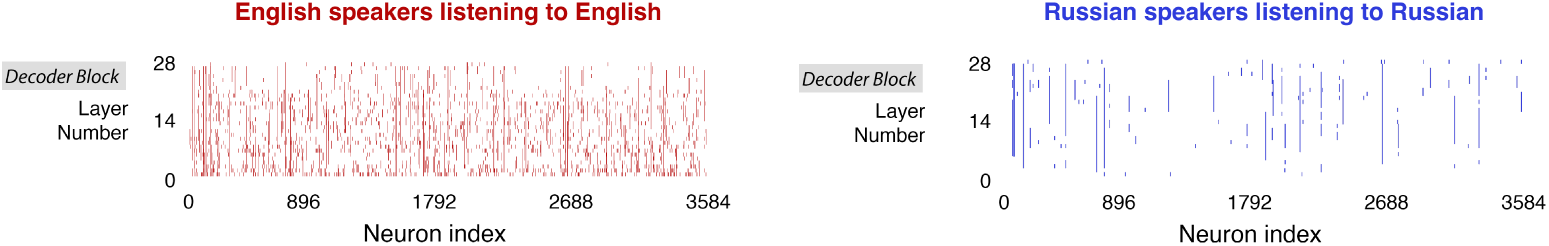

Conversely, the alignment pattern in the encoder block exhibits a distinct bipartite repetition, duplicating at the midpoint (neuron index 640):

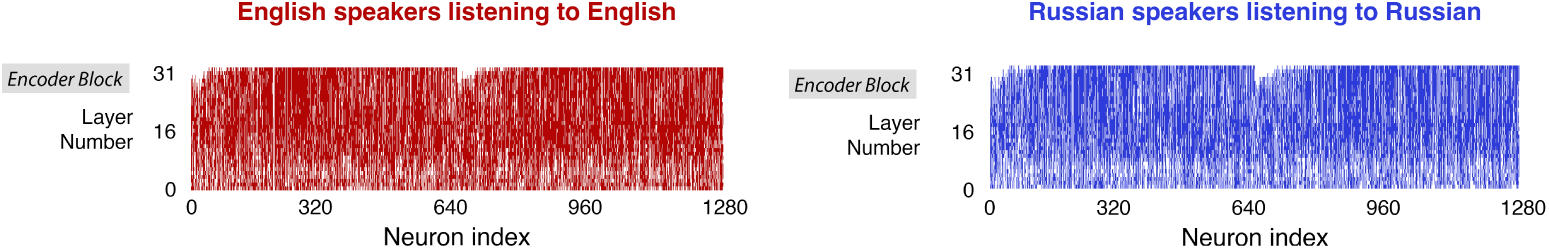

This structural repetition appears in both English and Russian native speakers processing their respective native languages. This topographic organization is a consequence of the encoder’s sine and cosine positional frequency embeddings, which are strictly segregated and mapped to the first and second halves of the neuron indices, respectively.

## Appendix SI2 Sustained neural alignment and the transformer residual stream

In the index-ordered alignment maps (Fig. 1B), the encoder block shows a bipartite repeat about index 640. This longitudinal persistence is a consequence of the transformer architecture’s residual stream [2]. Unlike strictly sequential feed-forward networks where representations are entirely reconstructed at each depth, the transformer architecture employs additive skip connections. These connections act as an identity bypass, allowing latent information to propagate forward through the network depth without being actively transformed by the intermediate attention or multi-layer perceptron (MLP) blocks.

Consequently, the specific neuron indices of the residual stream maintain a functional consistency across network depth. Once a useful linguistic or acoustic feature is written into a specific index and achieves significant cortical alignment at a given layer, the additive nature of the residual stream often ensures that this feature is retained—or only slowly morphed—as it propagates forward. This results in the observed phenomenon where a single index maintains its significant MEG alignment across numerous subsequent layers until the network actively overwrites that dimensional space.

While mapping the precise lifecycle of these residual features offers a promising avenue for tracking how specific linguistic computations emerge, stabilize, and decay across the cortical hierarchy, such a longitudinal analysis falls outside the scope of the present study. We note this topographical phenomenon here simply to contextualize the sustained vertical banding observed in the alignment matrices.

## Appendix SI3 Spatial topography of aLLM alignments in cortical source space

While the primary focus of this manuscript is the temporal and computational divergence point of speech processing, we also tested the aLLM-MEG alignments in cortical source space to verify their spatial distribution. To visualize this, for each cortical source, we identified the specific decoder layer containing the single highest-matching artificial neuron. The anatomical topography of these peak alignments, color-coded by layer depth, is presented in Supplementary Fig. SI1.

**Fig. SI1.**
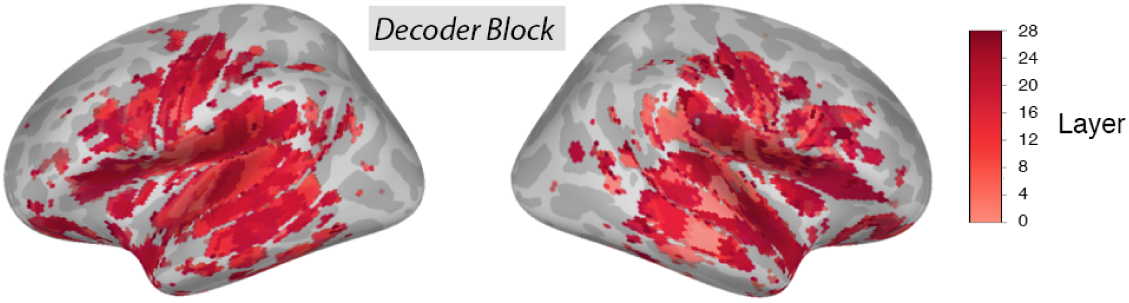
Source localization of Qwen2.5-Omni aLLM-brain alignment (highest-matching decoder layers.

Significant cortical tracking is distributed bilaterally across classical lateral temporal regions, including the auditory cortex, superior temporal gyrus (STG), and middle temporal gyrus (MTG). However, plotting the layer depth of these peak matches reveals a pronounced functional asymmetry. While both hemispheres exhibit widespread tracking, peak alignments in the right hemisphere are predominantly driven by the shallower layers of the decoder block. Conversely, the left hemisphere is heavily dominated by alignments to significantly deeper layers (indicated by darker colouration). Because this spatial projection assigns each source to its single best-matching unit, this pattern indicates that while foundational processing is bilateral, the higher-order lexico-semantic representations constructed in the deep layers are robustly left-lateralized, successfully outcompeting lower-level feature matches in the left hemisphere.

**Table SI1.** Comparison of linear and quadratic models on all datasets.

**Fig. SI1 | Source localization of Qwen2.5-Omni aLLM-brain alignment (highest-matching decoder layers).**
|  |  | Comprehension datasets |  |  |  | Non-comprehension dataset |  |
| --- | --- | --- | --- | --- | --- | --- | --- |
|  |  | English MEG [3] |  | Russian MEG [3] |  | Russ. lang., Eng. participants |  |
|  |  | Encoder fit | Decoder fit | Encoder fit | Decoder fit | Encoder fit | Decoder fit |
| **Monotonic trend** |  |  |  |  |  |  |  |
| | Spearman's $\rho$ | 0.927 | 0.913 | 0.880 | 0.597 | 0.597 | -0.309 |
| | $p$ | $2.76 \times 10^{-14}$ | $5.22 \times 10^{-12}$ | $3.34 \times 10^{-11}$ | $8.05 \times 10^{-4}$ | $6.13 \times 10^{-16}$ | n.s. ($3.85 \times 10^{-1}$) |
| **Linear model** |  |  |  |  |  |  |  |
|  | Fig. | SI2C | SI2A | SI2G | SI2E | SI2K | SI2I |
| | $R^2$ | 86.4% | 85.57% | 82.71% | 22.23% | 91.17% | 14.9% |
| | $p$ | $1.58 \times 10^{-14}$ | $5.13 \times 10^{-13}$ | $5.82 \times 10^{-13}$ | $1.13 \times 10^{-2}$ | $2.35 \times 10^{-17}$ | n.s. ($2.71 \times 10^{-1}$) |
|  | BIC | 196.96 | 266.59 | 162.55 | 285.04 | 183.45 | 104.35 |
| **Quadratic model** |  |  |  |  |  |  |  |
|  | Fig. | SI2D | SI2B | SI2H | SI2F | SI2L | SI2J |
| | $R^2$ | 85.8% | 91.28% | 81.8% | 32.22% | 82.0% | 2.3% |
| | $p$ | $3.02 \times 10^{-14}$ | $9.17 \times 10^{-16}$ | $1.32 \times 10^{-12}$ | $1.63 \times 10^{-3}$ | $1.12 \times 10^{-12}$ | n.s. ($6.76 \times 10^{-1}$) |
|  | BIC | 198.34 | 251.98 | 164.28 | 281.19 | 206.32 | 105.73 |
| **Comparison** |  |  |  |  |  |  |  |
| | $\Delta\text{BIC}(\text{linear} > \text{quadratic})$ | 1.38 | -14.60 | 1.73 | -3.85 | 22.88 | 1.38 |
|  | Preferred model | Linear | Quadratic | Linear | Quadratic | Linear | Linear |
|  | Strength | Weak | Very strong | Weak | Positive | Very strong | Weak |

**Table SI2.**
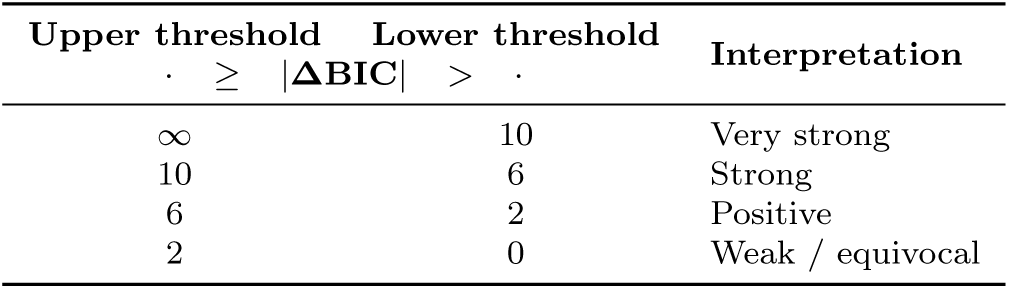
Rule-of-thumb thresholds for interpreting ΔBIC values.

## Appendix SI4 Comparison of linear and quadratic latency trends

Per-dataset latency fits and model comparisons are shown below; see Methods for the modelling procedure.

To quantify the size of the difference, we used the rule-of-thumb thresholds for |ΔBIC| values shown in Table SI2 (see [1] p. 139).

The results of fitting linear and quadratic models to the trends is show in Table SI1. In addition, the predictions from each model on each dataset are shown in Figure SI2. In all cases, the quadratic model was preferred for the decoder. For the English data, the quadratic model was strongly (ΔBIC *<* −6.0) or very strongly (ΔBIC *<* −10.0) preferred. For the Russian data the quadratic model was positively preferred (ΔBIC *<* −2.0).

**Fig. SI2.**
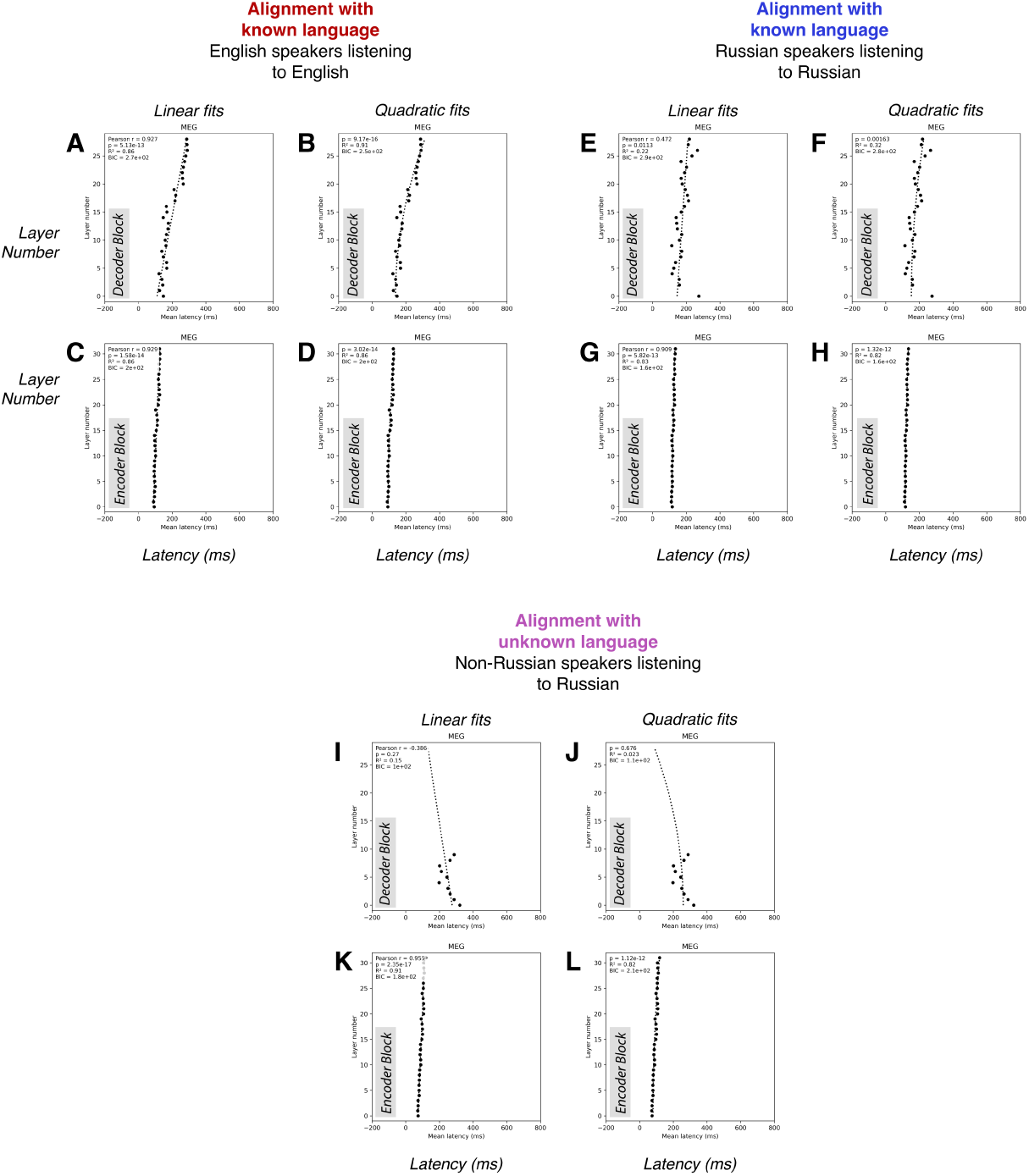
Linear and quadratic models for average latency as a function of layer. Black dots show the average latency for alignment for each layer (for layers with *≥* 4 significantly aligned neurons). The dotted line shows the trend. The left of each pair of graphs shows the linear model; the right shows the quadratic model. The top row shows decoder models; the bottom row shows encoder models. **(A–D)**: English stimulus, English participants. **(E–H)**: Russian stimulus, Russian participants. **(I–L)**: Russian stimulus, English participants.

Over all datasets and modalities with significant neuron fits, there is substantial evidence that latency increases quadratically with the depth of the neuron in the decoder block.

## References

[1] Friederici, A.D.: The cortical language circuit: from auditory perception to sentence comprehension. Trends in cognitive sciences 16(5), 262–268 (2012)

[2] Davis, M.H., Johnsrude, I.S.: Hierarchical processing in spoken language comprehension. Journal of Neuroscience 23(8), 3423–3431 (2003)

[3] Howard, M.F., Poeppel, D.: Discrimination of speech stimuli based on neuronal response phase patterns depends on acoustics but not comprehension. Journal of Neurophysiology 104(5), 2500–2511 (2010) 10.1152/jn.00251.2010

[4] Zoefel, B., VanRullen, R.: EEG oscillations entrain their phase to high-level features of speech sound. NeuroImage 124, 16–23 (2016) 10.1016/j.neuroimage.2015.08.054

[5] Kutas, M., Federmeier, K.D.: Electrophysiology reveals semantic memory use in language comprehension. Trends in cognitive sciences 4(12), 463–470 (2000)

[6] Petit, S., Badcock, N.A., Grootswagers, T., Rich, A.N., Brock, J., Nickels, L., Moerel, D., Dermody, N., Yau, S., Schmidt, E., Woolgar, A.: Toward an individualized neural assessment of receptive language in children. Journal of Speech, Language, and Hearing Research 63(7), 2361–2385 (2020)

[7] Oota, S.R., Pahwa, K., Marreddy, M., Gupta, M., Raju, B.S.: Neural architecture of speech. In: ICASSP 2023-2023 IEEE International Conference on Acoustics, Speech and Signal Processing (ICASSP), pp. 1–5 (2023). IEEE

[8] Tuckute, G., Sathe, A., Srikant, S., Taliaferro, M., Wang, M., Schrimpf, M., Kay, K., Fedorenko, E.: Driving and suppressing the human language network using large language models. Nature Human Behaviour 8(3), 544–561 (2024)

[9] Millet, J., Caucheteux, C., Boubenec, Y., Gramfort, A., Dunbar, E., Pallier, C., King, J.-R., et al.: Toward a realistic model of speech processing in the brain with self-supervised learning. Advances in Neural Information Processing Systems 35, 33428–33443 (2022)

[10] Tuckute, G., Feather, J., Boebinger, D., McDermott, J.H.: Many but not all deep neural network audio models capture brain responses and exhibit correspondence between model stages and brain regions. Plos Biology 21(12), 3002366 (2023)

[11] Li, Y., Anumanchipalli, G.K., Mohamed, A., Chen, P., Carney, L.H., Lu, J., Wu, J., Chang, E.F.: Dissecting neural computations in the human auditory pathway using deep neural networks for speech. Nature Neuroscience 26(12), 2213–2225 (2023)

[12] Caucheteux, C., Gramfort, A., King, J.-R.: Evidence of a predictive coding hierarchy in the human brain listening to speech. Nature human behaviour 7(3), 430–441 (2023)

[13] Schrimpf, M., Blank, I.A., Tuckute, G., Kauf, C., Hosseini, E.A., Kanwisher, N., Tenenbaum, J.B., Fedorenko, E.: The neural architecture of language: Integrative modeling converges on predictive processing. Proceedings of the National Academy of Sciences 118(45), 2105646118 (2021) 10.1073/pnas.2105646118

[14] Gwilliams, L., Marantz, A., Poeppel, D., King, J.-R.: Hierarchical dynamic coding coordinates speech comprehension in the brain. bioRxiv (2024)

[15] Hadidi, N., Feghhi, E., Song, B.H., Blank, I.A., Kao, J.C.: Spurious alignment between large language models and brains can emerge from non-robust methods and overlooked confounds. Nature Communications (2026) 10.1038/s41467-026-72253-7

[16] Ding, N., Melloni, L., Zhang, H., Tian, X., Poeppel, D.: Cortical tracking of hierarchical linguistic structures in connected speech. Nature Neuroscience 19, 158–164 (2016)

[17] Bhaya-Grossman, I., Leonard, M.K., Zhang, Y., Gwilliams, L., Johnson, K., Lu, J., Chang, E.F.: Shared and language-specific phonological processing in the human temporal lobe. Nature 649(8095), 140–151 (2026)

[18] Schnakers, C., Vanhaudenhuyse, A., Giacino, J., Ventura, M., Boly, M., Majerus, S., Moonen, G., Laureys, S.: Diagnostic accuracy of the vegetative and minimally conscious state: clinical consensus versus standardized neurobehavioral assessment. BMC Neurology 9(1), 35 (2009) 10.1186/1471-2377-9-35

[19] Gui, P., Jiang, Y., Zang, D., Qi, Z., Tan, J., Tanigawa, H., Jiang, J., Wen, Y., Xu, L., Zhao, J., Mao, Y., Poo, M.-M., Ding, N., Dehaene, S., Wu, X., Wang, L.: Assessing the depth of language processing in patients with disorders of consciousness. Nature Neuroscience 23(6), 761–770 (2020) 10.1038/s41593-020-0639-1

[20] Xu, J., Guo, Z., He, J., Hu, H., He, T., Bai, S., Chen, K., Wang, J., Fan, Y., Dang, K., Zhang, B., Wang, X., Chu, Y., Lin, J.: Qwen2.5-Omni Technical Report (2025). https://arxiv.org/abs/2503.20215

[21] Glasberg, B.R., Moore, B.C.J.: A model of loudness applicable to time-varying sounds. Journal of the Audio Engineering Society 50(5), 331–342 (2002)

[22] Peelle, J.E., Johnsrude, I., Davis, M.H.: Hierarchical processing for speech in human auditory cortex and beyond. Frontiers in human neuroscience 4, 1735 (2010)

[23] Peelle, J.E., Gross, J., Davis, M.H.: Phase-locked responses to speech in human auditory cortex are enhanced during comprehension. Cerebral cortex 23(6), 1378–1387 (2013)

[24] Peña, M., Melloni, L.: Brain oscillations during spoken sentence processing. Journal of Cognitive Neuroscience 24(5), 1149–1164 (2012) 10.1162/jocn_a_00144

[25] Goldstein, A., Zada, Z., Buchnik, E., Schain, M., Price, A., Aubrey, B., Nastase, S.A., Feder, A., Emanuel, D., Cohen, A., et al.: Shared computational principles for language processing in humans and deep language models. Nature neuroscience 25(3), 369–380 (2022)

[26] Paulo, G., Mallen, A., Juang, C., Belrose, N.: Automatically Interpreting Millions of Features in Large Language Models (2025). https://arxiv.org/abs/2410.13928

[27] Bills, S., Cammarata, N., Mossing, D., Tillman, H., Gao, L., Goh, G., Sutskever, I., Leike, J., Wu, J., Saunders, W.: Language models can explain neurons in language models (2023). https://openaipublic.blob.core.windows.net/neuron-explainer/paper/index.html

[28] Kriegeskorte, N., Douglas, P.K.: Cognitive computational neuroscience. Nature Neuroscience 21, 1148–1160 (2018) 10.1038/s41593-018-0210-5

[29] Yang, C., Parish, O., Klimovich-Gray, A., Wingfield, C., Marslen-Wilson, W.D., Zhang, C., Woolgar, A., Thwaites, A.: Kymata Soto language dataset: an electro-magnetoencephalographic dataset for natural speech processing. Scientific Data (2026)

[30] Taulu, S., Simola, J., Kajola, M.: Applications of the signal space separation method. IEEE transactions on signal processing 53(9), 3359–3372 (2005)

[31] Hämäläinen, M.S., Ilmoniemi, R.J.: Interpreting magnetic fields of the brain: minimum norm estimates. Medical & biological engineering & computing 32(1), 35–42 (1994)

[32] McAuliffe, M., Socolof, M., Mihuc, S., Wagner, M., Sonderegger, M.: Montreal forced aligner: Trainable text-speech alignment using kaldi. In: Interspeech, vol. 2017, pp. 498–502 (2017)

[33] King, J.-R., Charton, F., Lopez-Paz, D., Oquab, M.: Back-to-back regression: Disentangling the influence of correlated factors from multivariate observations. NeuroImage 220, 117028 (2020)

[34] Thwaites, A., Glasberg, B.R., Nimmo-Smith, I., Marslen-Wilson, W.D., Moore, B.C.: Representation of instantaneous and short-term loudness in the human cortex. Frontiers in Neuroscience 10, 183 (2016)

[35] Thwaites, A., Schlittenlacher, J., Nimmo-Smith, I., Marslen-Wilson, W.D., Moore, B.C.: Tonotopic representation of loudness in the human cortex. Hearing research 344, 244–254 (2017)

[36] Thwaites, A., Zhang, C., Woolgar, A.: Information processing pathway maps — a scalable framework for mapping cortical processing. NeuroImage 317, 121345 (2025) 10.1016/j.neuroimage.2025.121345

[37] Raftery, A.E.: Bayesian model selection in social research. Sociological methodology, 111–163 (1995)

[38] Glasberg, B.R., Moore, B.C.J.: A model of loudness applicable to time-varying sounds. Journal of the Audio Engineering Society 50(5), 331–342 (2002)

[39] Kruskal, W.H., Wallis, W.A.: Use of ranks in one-criterion variance analysis. Journal of the American statistical Association 47(260), 583–621 (1952)

## Supplementary References

[1] Adrian E Raftery. Bayesian model selection in social research. Sociological methodology, pages 111–163, 1995.

[2] Ashish Vaswani, Noam Shazeer, Niki Parmar, Jakob Uszkoreit, Llion Jones, Aidan N Gomez, Lukasz Kaiser, and Illia Polosukhin. Attention is all you need. Advances in neural information processing systems, 30, 2017.

[3] ChenTianyi Yang, Oliver Parish, Anastasia Klimovich-Gray, Cai Wingfield, William D. Marslen-Wilson, Chao Zhang, Alexandra Woolgar, and Andrew Thwaites. Kymata Soto language dataset: an electro-magnetoencephalographic dataset for natural speech processing. Scientific Data, 2026.

